# Multilocus sex determination revealed in two populations of gynodioecious wild strawberry, *Fragaria vesca* subsp. *bracteata*

**DOI:** 10.1101/023713

**Authors:** T-L. Ashman, J.A. Tennessen, R. Dalton, R. Govindarajulu, M. Koski, A. Liston

**Author notes:** **Author for correspondence:** Tia-Lynn Ashman Department of Biological Sciences, 4249 Fifth Ave, Pittsburgh, PA 15260).

## Abstract

Gynodioecy, the coexistence of females and hermaphrodites, occurs in 20% of angiosperm families and often enables transitions between hermaphroditism and dioecy. Clarifying mechanisms of sex determination in gynodioecious species can thus illuminate sexual system evolution. Genetic determination of gynodioecy, however, can be complex and is not fully characterized in any wild species. We used targeted sequence capture to genetically map a novel nuclear contributor to male sterility in a self-pollinated hermaphrodite of *Fragaria vesca* subsp. *bracteata* from the southern portion of its range. To understand its interaction with another identified locus and possibly additional loci, we performed crosses within and between two populations separated by 2000 km, phenotyped the progeny and sequenced candidate markers at both sex-determining loci. The newly mapped locus contains a high density of pentatricopeptide repeat genes, a class commonly involved in restoration of fertility caused by cytoplasmic male sterility. Examination of all crosses revealed three unlinked epistatically interacting loci that determine sexual phenotype and vary in frequency between populations. *Fragaria vesca* subsp. *bracteata* represents the first wild gynodioecious species with genomic evidence of both cytoplasmic and nuclear genes in sex determination. We propose a model for the interactions between these loci and new hypotheses for the evolution of sex determining chromosomes in the subdioecious and dioecious *Fragaria.*

## INTRODUCTION

Diversity in sexual systems and sex determination is a hallmark of plants (Bachtrog et al. 2014; Renner 2014). In angiosperms in particular, this variety is indicative of the myriad ways unisexual individuals can evolve from combined sex phenotypes (hermaphrodites or cosexuals) (Diggle et al. 2011; Renner 2014). One canonical pathway to entirely separate sexes (dioecy) involves an intermediate sexual system known as gynodioecy (females and hermaphrodites) (Charlesworth and Charlesworth 1978) which is found in nearly 20% of families and 2% of genera of flowering plants (Dufaÿ et al. 2014). Gynodioecy, however, does not always lead to dioecy and may be a stable sexual system in its own right (Lewis 1941; Charlesworth and Charlesworth 1978; Dufay et al. 2007), or may transition back to hermaphroditism (Delph et al. 2007; Goldberg et al. unpublished). The likelihood of these transitions is influenced by the underlying genetics, among other factors such as mating system and frequency dependent selection (Ehlers and Bataillon 2007; Crossman and Charlesworth 2014).

Characterizing the mechanisms of sex determination in gynodioecious species can be key for understanding sexual system transitions (Charlesworth and Charlesworth 1978; Schultz 1994; Maurice and Fleming 1995; Bailey and Delph 2007), as well as for elucidating aspects of male and female reproductive developmental pathways (Wang et al 2013; Diggle et al 2011). Known sex determination systems can involve cytoplasmic and/or nuclear genes. In cyto-nuclear gynodioecy, mitochondrial male-sterility mutations (cytoplasmic male sterility, ‘CMS’) are counteracted by alleles at nuclear ‘restorer’ (*Rf*) loci (e.g., Dufay et al. 2009). Restorers are diverse, but most commonly are pentatricopeptide repeat (PPR) genes that act in dominant fashion so only one copy is required to restore pollen function (sterility at *Rf* loci is typically recessive) (reviewed by Chen and Liu 2014). Nuclear sex determination can result when CMS is fixed and only restorers segregate (Klaas and Olson 2006), or when nuclear mutations cause loss-of-function (LOF) in the pollen development pathway irrespective of cytoplasmic genes (Bailey and Delph 2007). Finally, in some species both cytoplasmic and nuclear mutations segregate within populations and there can be more than one nuclear locus (e.g., Van Damme et al. 2004; Garraud et al. 2011). Reversions to hermaphroditism can occur within populations when CMS or nuclear LOF genes are lost or when both CMS and restorers become fixed (e.g., *Arabidopsis thaliana,* Gobron et al. 2013). As a consequence, disparate populations of a widespread species that have been exposed to dissimilar selection regimes or differentially subjected to drift (population size or founder events) can be differentiated for sex determination mechanisms. Such variation has been revealed in species with cyto-nuclear gynodioecy (e.g., *Lobelia siphiltica,* Dudle et al. 2001), but also can be seen in species with nuclear sex determination or sex chromosomes (e.g., Dufresnes et al. 2014). In these cases, transitions in sex determining regions (or chromosomes) are thought to be driven by additional selective forces, such as sexually antagonistic selection, meiotic drivers and genetic load (e.g., Blaser et al. 2014; Ubeda et al. 2015).

Model organisms and agriculturally important taxa have been instrumental in characterizing sex determination pathways and have revealed that transitions between sexual systems can involve new genes, new alleles or entirely novel means of sex determination (e.g., Vicoso and Bachtrog 2013; Akagi et al. 2014). In fact, the same mechanism can be regulated by different genes, and thus developmental networks can evolve rapidly (Cui et al. 2012; Wang et al. 2013). Understanding sex determination in wild plants is particularly valuable because it is in these species that data can be linked to ecological processes that are prevailing drivers of sexual system variation (Frank 1989; Jacobs and Wade 2003). Gynodioecious species are often closely related to hermaphroditic or dioecious ones (Dufaÿ et al. 2014), so characterization of their sex determination can also provide insight into genetic, developmental and evolutionary dynamics of sexual systems (Spigler and Ashman 2012; Russell and Pannell 2015).

We explored variation in sex determination in the widespread North American diploid strawberry *F. vesca* subsp. *bracteata* for two main reasons. First, it is gynodioecious and the maternal donor to the sexually dimorphic octoploid *Fragaria* (dioecious *F. chiloensis* and subdioecious *F. virginiana*) (Njuguna et al. 2013; Tennessen et al. 2014; Govindarajulu et al. 2015) which were hybridized to produce the cultivated hermaphroditic *F.* ×*ananassa*. As such it represents both a tractable and evolutionarily appropriate genetic model for teasing apart sex differentiation in this crop and its wild relatives (Liston et al. 2014). Second, several lines of evidence allude to variation in sex determining regions within the species. Specifically, although previous genetic mapping of sex determination in a northern population of *F. vesca* subsp. *bracteata* identified a nuclear locus with a dominant allele for male sterility (Tennessen et al. 2013), there is equivocal evidence for a fitness advantage sufficient to maintain females in this population under nuclear determination alone (Li et al. 2012; Dalton et al. 2013). Moreover, recent phylogeographic studies revealed genetic differentiation across the range of *F. vesca* subsp. *bracteata*, with one combination of chlorotype and mitotype more closely related to octoploids than other(s) (Njuguna et al. 2013; Govindarajulu et al. 2015; Stanley et al. 2015). Geographic differentiation was partly due to variation in a novel mitochondrial open reading frame (ORF) with sequence similarity to known CMS genes, although a sterilizing function has not been confirmed (Govindarjulu et al. 2015; Stanley et al 2015). These points combined with the fact that the location of the sex determining region of *F. vesca* subsp. *bracteata* identified in Tennessen et al. (2013) was on a chromosome that is different from either of the chromosomal locations of the sex determining region in the two octoploid congeners (i.e., chromosome 4 (LG4) *vs.* a chromosome in homeologous group VI (LG VI-Av or VI-B2), Goldberg et al. 2010; Spigler et al. 2011; Tennessen et al. 2014) raises the possibility of additional genetic contributors to sex determination in other populations of the diploid species.

We used targeted-sequence capture (Tennessen et al 2013; 2014) to efficiently fine map the sex determination region using a self-pollinated hermaphrodite plant from a population at the southeastern edge of *F. vesca* subsp. *bracteata*’s range within the USA. From this we identified a second locus affecting sexual phenotype on a different chromosome (LG6) than that previously identified from a female collected in the northwestern portion of the range (LG4; Tennessen et al. 2013). We conducted intra- and inter-population crosses. In addition to phenotyping the progeny, we sequenced informative SNPs near the target sex-determining regions to determine whether the second locus interacted with the locus previously identified. In doing, so we reveal interactions of at least two nuclear loci and population variation in sex determination. From this we propose a novel mechanistic pathway for sex determination in the diploid species, and extend it as a hypothesis for sex determination in the octoploid species that descended from the diploid species studied here.

## MATERIALS AND METHODS

### Study system

*Fragaria vesca* subsp. *bracteata* (Rosaceae) is a diploid (2n = 2× = 14) North American wild strawberry that inhabits moderately damp soils of shady forest edges, meadows, and riverbanks. Its range extends from British Columbia to southern Mexico and from the Pacific coast to the Rocky Mountains and Sierra Madre (Staudt 1999; Stanley et al. 2015). Variation in cytoplasmic haplotypes has been revealed across this range with western populations dominated by one chlorotype and those from the Rockies and Sierra Madre by another with a zone of admixture in between (Njuguna et al. 2013; Stanley et al. 2015). In addition, there is finer scale and more complex pattern of mitochondrial variation (Stanley et al. 2015).

The species has a gynodioecious sexual system (Ahmadi and Bringhurst 1989; Li et al. 2012) and populations vary in the frequency of females from 0 (no females) to 46% female (Stanley et al. 2015). Sex types produce similar numbers of flowers, seeds and plantlets on stolons (Li et al. 2012). Hermaphrodites are self-compatible, highly selfing and morphologically distinguishable from females by their slightly longer stamens and the presence of viable pollen grains in their anthers (Li et al. 2012; Dalton et al. 2013).

The only known nuclear locus that affects sex expression was identified from a cross between a female and a hermaphrodite from Oregon, USA (OR-MRD; Tennessen et al. 2013). The dominant allele for male sterility resides in a 338-kb region of chromosome 4 that is predicted to house 57 genes, although none in protein families known to control male sterility were found based on the reference genome *F. vesca* Hawaii4 annotations (Tennessen et al. 2013). However, Govindarajulu et al. (2015) identified a mitochondrial ORF (*atp8-orf225*) with molecular characteristics similar to known CMS genes. It is present and variable in *F. vesca* subsp. *bracteata* as well as other taxa across the genus *Fragaria* (Govindarajulu et al. 2015; Stanley et al. 2015), but its functional role has yet to be confirmed.

### Study materials

*Fragaria vesca* subsp. *bracteata* from two populations separated by 2000 km were the focus of the current study. NM-LNF is a high elevation ‘sky island’ population in the southern portion of the range (in the Lincoln National Forest, New Mexico, altitude 2664 m; N 32° 58′ 6.1′′, W 105° 44′ 44.7′′). OR-MRD is on the highest peak in the Oregon Coast Range (Marys Peak in the Siuslaw National Forest, Oregon, altitude 521 m; N 44.8° 29′ 18.4′′, W 123° 32′ 14.7′′). The latter population is also within the zone of cytoplasmic introgression seen in *F. vesca* subsp. *bracteata* between the Coast Range / Cascades and Northern Rocky Mountains (Staudt 1999; Stanley et al. 2015). The frequency of females in OR-MRD is 30% and in NM-LNF is 46% (Stanley et al. 2015).

Geography, flower and fruit morphology, and sexual system indicate both populations are *F. vesca* subsp. *bracteata* (Staudt 1999; T-L. Ashman, unpublished). Likewise phylogenomic analysis shows these populations form a clade for the majority of their chromosomes (Tennessen et al. 2014) and are always separate from *F. vesca* subsp. *vesca* Hawaii 4 (source of the published genome sequence, Shulaev et al, 2010) which itself forms a clade with *F. vesca* subsp. *americana* (Njuguna et al 2013). These two populations, however, differed in their cytotypes as is common in *F. vesca* subsp. *bracteata*: OR-MRD has one chlorotype (2) and two mitotypes (90% C and 10% B), whereas NM-LNF is fixed for different chlorotype and mitotype (1 and F, respectively; Stanley et al. 2015).

### Crosses and sexual phenotyping

Plants were collected from the wild and maintained in the greenhouse at the University of Pittsburgh. Three hermaphrodites (OR-MRD61, OR-MRD45, OR-MRD93) and three females (OR-MRD30 [used in Tennessen et al. 2013], OR-MRD27, OR-MRD90) from OR-MRD were used as parents. From NM-LNF, three hermaphrodites (NM-LNF23 [used in Tennessen et al. 2014], NM-LNF25, and NM-LNF14) and three females (NM-LNF2, NM-LNF4, NM-LNF26) were used as parents. Cytotypes of the parents reflected their abundance in wild populations (Stanley et al. 2015) and are given in Table S1. The three *atp8-orf225* variants translate to different proteins, although their function is unknown (Stanley et al. 2015).

All females were crossed with pollen from both inter- and intra-population hermaphrodites, and all hermaphrodites were self-pollinated. In addition, crosses between the hermaphrodites were performed within and between populations. Crosses were performed by hand-pollinating 1-4 flowers per dam with pollen collected from single sires. When hermaphrodites served as dams their flowers were emasculated prior to pollination. Seeds were harvested and stored at -20° C until sowing. Some crosses produced few seeds (and/or few germinated), notably hermaphrodite by hermaphrodite interpopulation crosses involving NM-LNF as the dam and OR-MRD as the sire, and some hermaphrodite by hermaphrodite within population crosses (see results Table 1). We sowed an average of 31 seeds (range 1-56) per family into 98 well trays in a mix of Fafard #4, sand, and Sunshine Germination Mix. Germination was, on average, high (75%), although some specific crosses performed poorly (see results). Approximately one month after germination we transplanted progeny into 150cc pots of soil (two parts Fafard #4 to 1 part sand) and randomized them on benches in a greenhouse where we maintained conditions of 14-15 hr days 21°/15.5° C (day/night) temperatures.

**Table 1.**
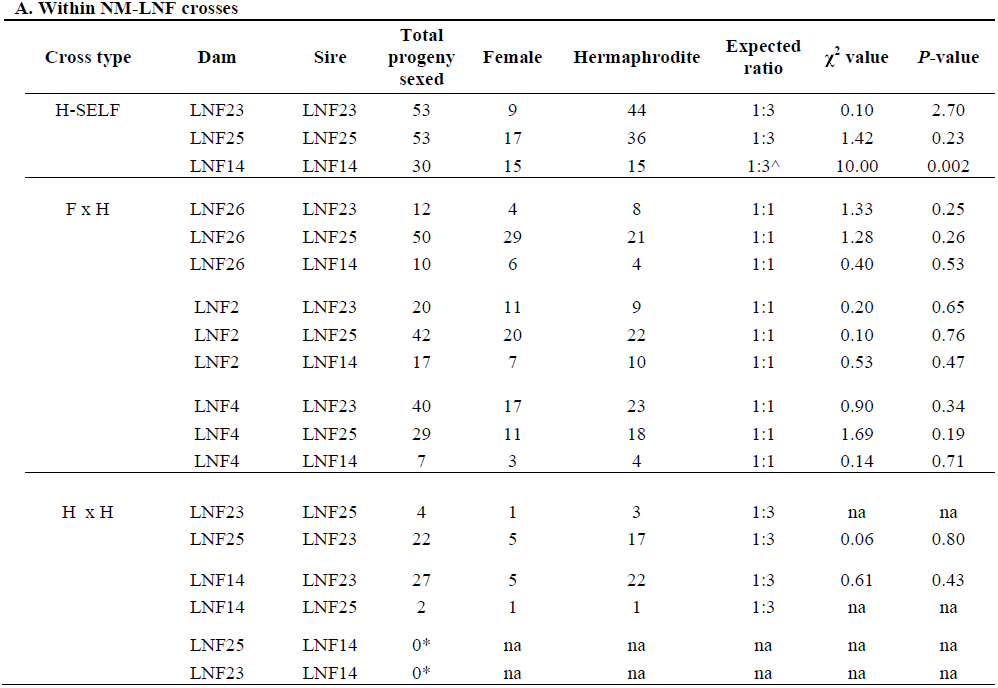
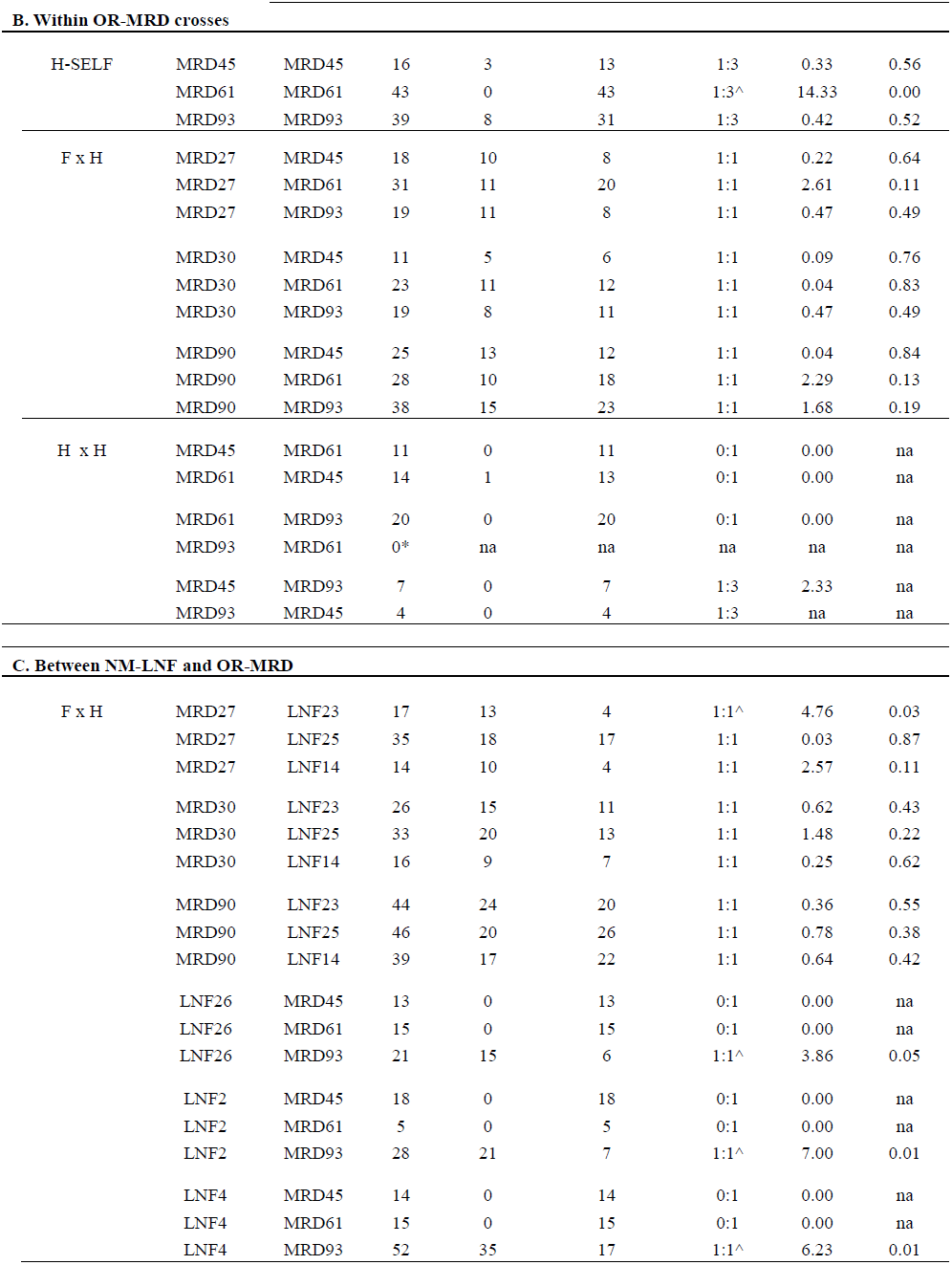
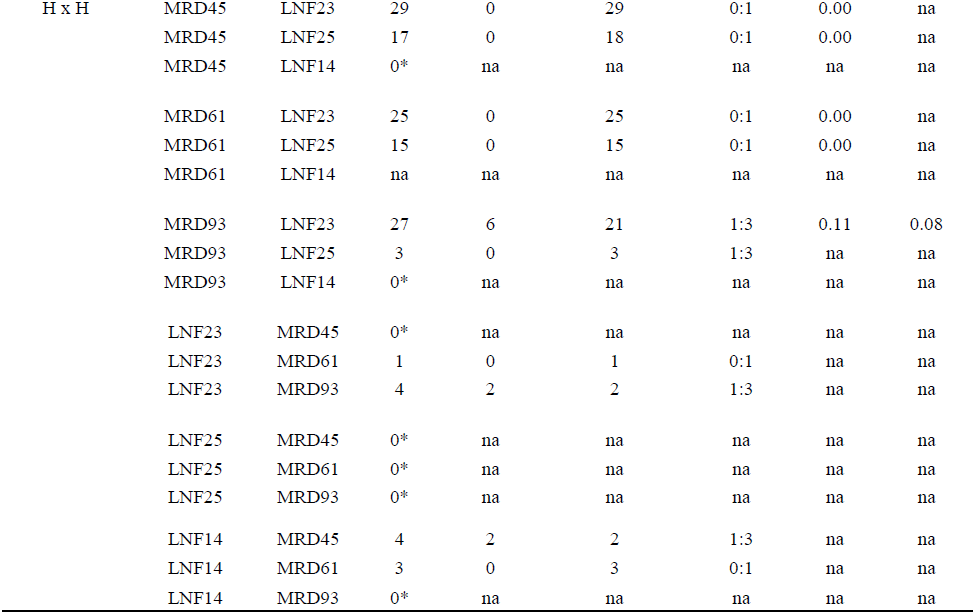
Phenotypic sex ratios from intra-(within population) and inter-population (between populations) crosses of gynodioeicous *Fragaria vesca* subsp. *bracteata*. Three types of crosses are presented: selfed hermaphrodites (H-self), female dam by hermaphrodite sire (F × H) and hermaphrodite dam by hermaphrodite sire (H × H) from within (A) N-LNF or (B) OR-MRD populations or between NM-LNF and OR-MRD populations (C). For each cross the dam, sire and total number of progeny scored for sexual phenotype are given, as well as the number of female and hermaphrodite progeny, the predicted sex ratio based on Table 3 genotypes of the parents. χ^2^ statistics and *P* values from χ^2^ goodness of fit tests. *Crosses performed yielded no seed or none germinated. ^Alternative hypothesis for expected sex ratio tested and reported in text. Na: Too few progeny (family size <10) to conduct a statistical test.

To induce flowering we subjected plants to a cold treatment (8 hr days at 8°C/16hr dark at 4°C) in a growth chamber for three to four weeks, and then returned them to the greenhouse where they received 50ppm 10:30:20 N:P:K fertilizer (Peter’s Professional(®) Bloom Booster) weekly. This was repeated up to five times to maximize flowering. A high rate of flowering was achieved (on average 86% progeny per family). Upon flowering we scored male function for at least two flowers per plant. Given the weak sexual dimorphism in this species (Li et al 2012), we recorded both whether or not the anthers were shedding pollen in the greenhouse and whether viable pollen grains were produced. We confirmed pollen production and viability microscopically by fixing anthers with Alexander’s stain (Kearns and Inouye 1993), and later observing them under a light microscope. Plants with no viable pollen production were scored as sterile whereas those with viable pollen production were scored as fertile. In total, 1364 plants flowered and the number of individuals phenotyped per family ranged from 1 to 53 (mean = 21). Progeny sex ratios were scored from flowering plants in each cross and for family size >10 sex ratio was tested for deviations from expected (see below) using χ^2^ tests.

### Genetic and genomic analysis

To identify the sex-determining locus in NM-LNF we used the linkage map published in Tennessen et al. (2014). This map was created using OneMap (Margarido et al. 2007) and the 1825 polymorphic sites identified from the target captured sequences of 41 progeny of self-pollinated NM-LNF23 (Tennessen et al. 2014). This map had the expected seven linkage groups spanning a total of 326 cM and is described in full in Tennessen et al (2014) where it was used to produce a new *Fragaria vesca* genome assembly (‘Fvb’). Based on past experience (Tennessen et al. 2013) this sample size is sufficient to identify genomic regions of major effect on male function. We coded male sterility as a recessive Mendelian locus (e.g., sterile *rr* and fertile *R-*; justified because a hermaphroditic parent could not harbor a dominant male sterility allele), and mapped it along with the targeted capture genotypes using OneMap (Margarido et al. 2007) to determine its position on the linkage map. From this map we identified a candidate region in coupling with male sterility on LG6 (see results).

Once the male function region was identified, we designed primers for PCR amplification and Sanger sequencing of three SNPs from target capture sequences on LG6: one just upstream (Fvb6_34763k), one within (Fvb6_35142k), and one downstream of the male sterility region (Fvb6_36607k) (Table S2; Figure 2). Primers were designed with Primer-BLAST (http://www.ncbi.nlm.nih.gov/tools/primer-blast/) and positioned to amplify additional nearby SNPs segregating in OR-MRD30, as well as the NM-LNF23 SNPs. To genotype the region housing the sex expression locus on LG4, we used primers described previously using the *F. vesca* Hawaii 4 assembly but that were re-aligned to the new assembly ‘Fvb’ (Fvb4_30092k) (Tennessen et al. 2014; Table S2).

Sequencing of targeted sex determining regions on LG4 and LG6 was performed on seven parents and 301 progeny from 13 select crosses. The crosses were selected based on the availability of segregating informative SNPs in the targeted regions in the parents (Table 2 I) and sexual phenotypes of their progeny. We sequenced progeny from three crosses for LG4 markers (OR-MRD30×NM-LNF23, OR-MRD30×NM-LNF14, and OR-MRD90×NM-LNF23) and progeny from ten crosses for LG6 markers (OR-MRD93self, NM-LNF14self, NM-LNF2× NM-LNF23, NM-LNF2×OR-MRD93, OR-MRD30×NM-LNF23, OR-MRD30×NM-LNF14, OR-MRD90×NM-LNF23, OR-MRD93×NM-LNF23, OR-MRD61×NM-LNF23, and OR-MRD61×OR-MRD93).

**Table 2.**
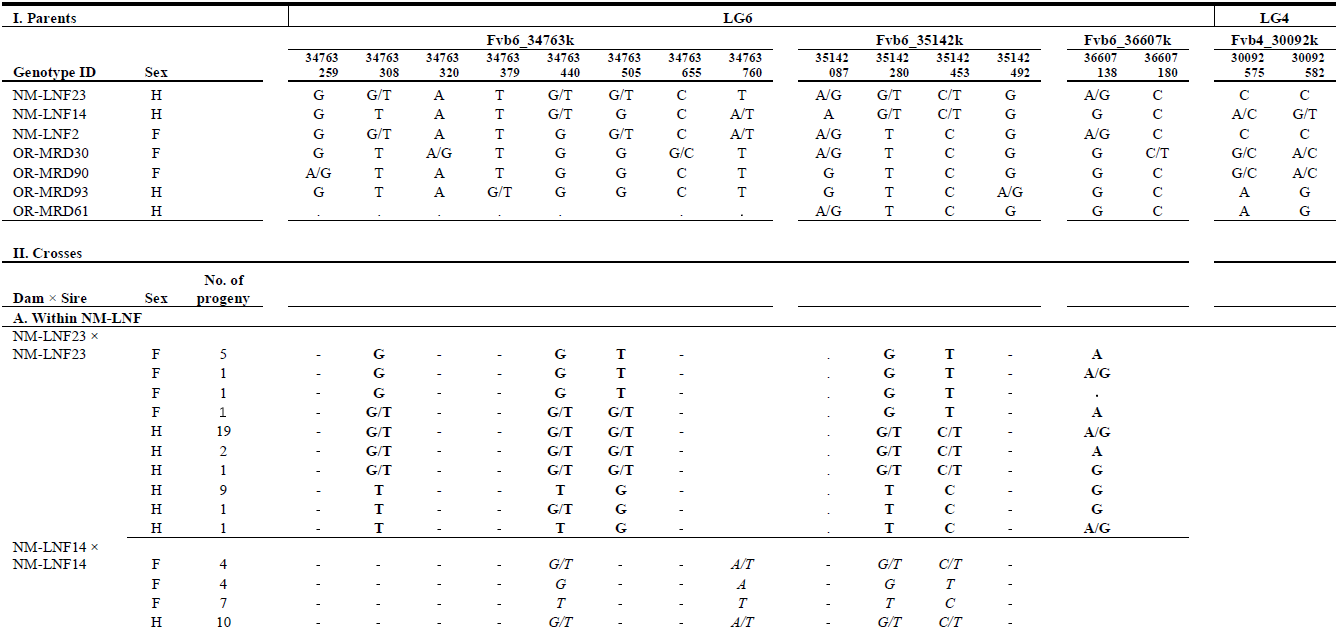
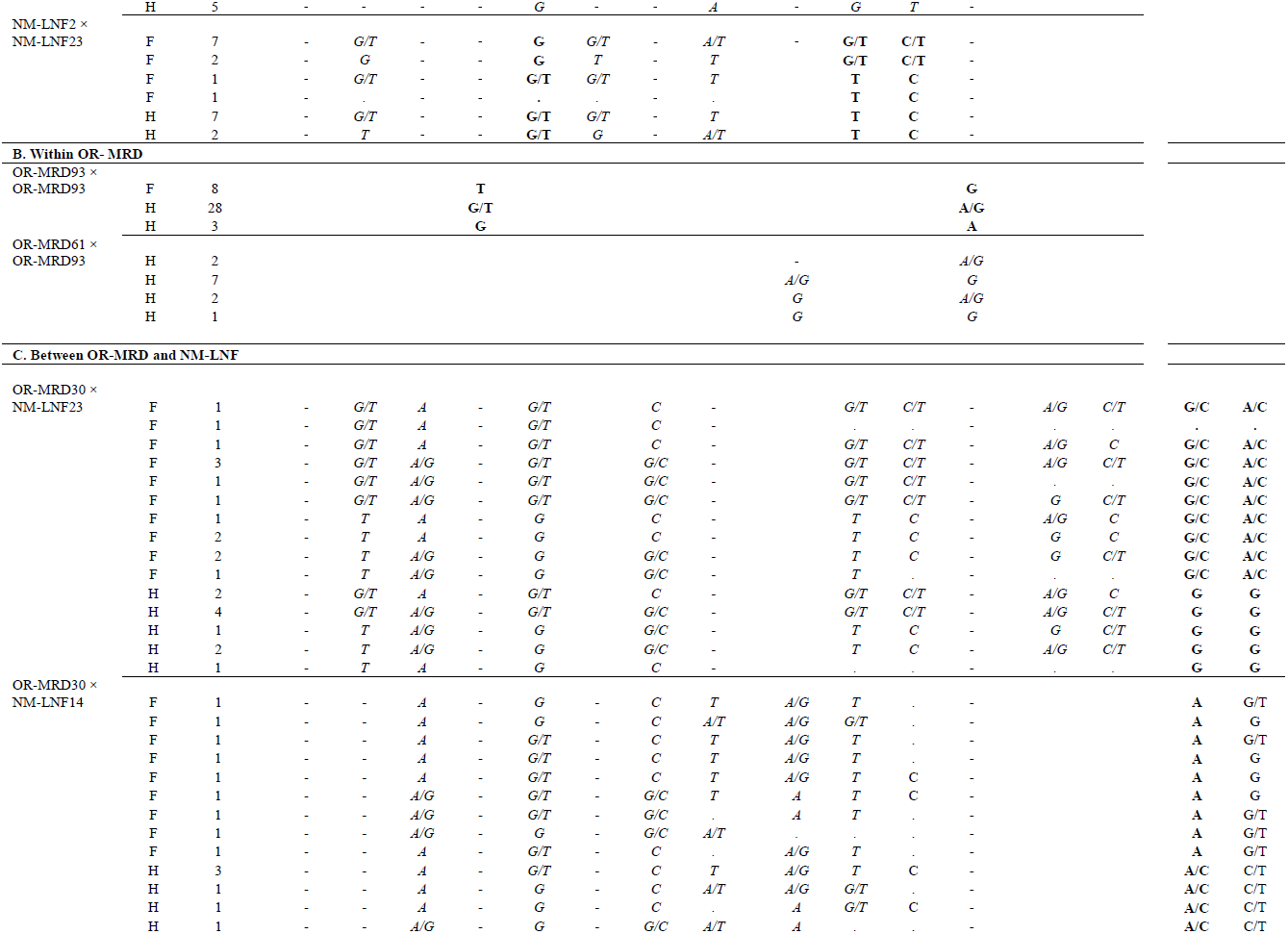
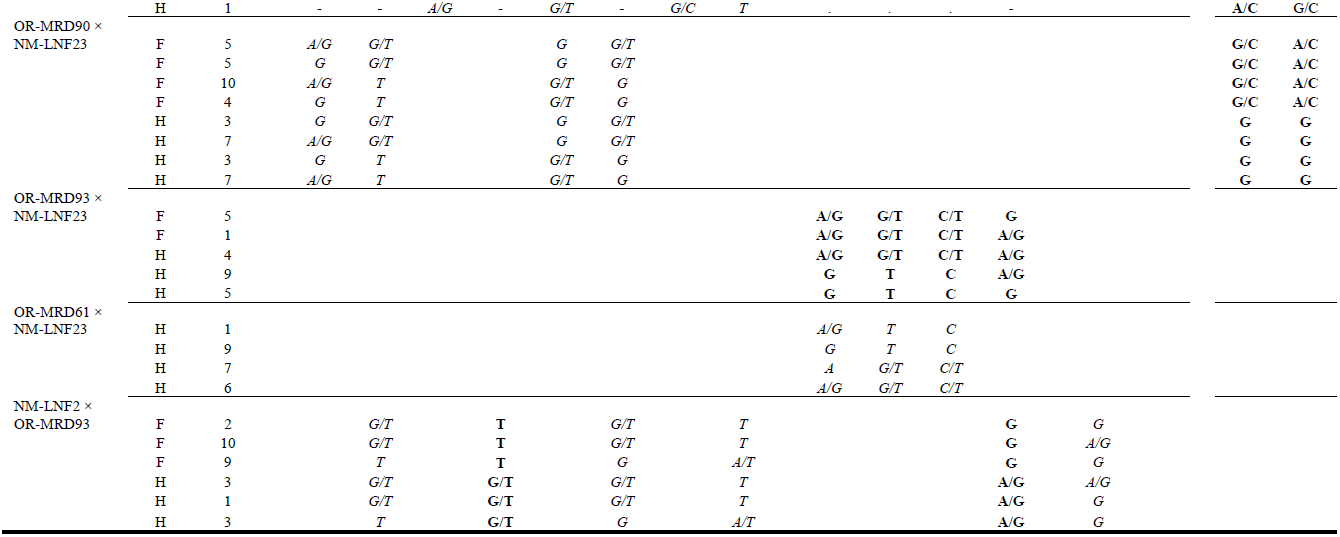
Single nucleotide polymorphic sites segregating at the putative sex determining regions on linkage groups 6 and 4 (LG6 and LG4) (see Figure 2AB) in two populations (OR-MRD, NM-LNF) of gynodioecious *Fragaria vesca* subsp. *bracteata*. Sexual phenotypes (F=female, H=hermaphrodite) and genotypes for parents (I) and progeny (II) following intra-(A, B) and inter-population (C) crosses. Genotypes in progeny are only shown for screened positions that were polymorphic in the parents. Dashes denote non-segregating SNPs, “.” indicate missing data, whereas empty cells reflect positions that were not sequenced. SNPs in bold are correlated with sex, while SNPs in italics are not.

For each cross we extracted DNA from 100-150 mg of fresh or 20-30 mg of silica-dried leaf tissue per plant. We used a CTAB DNA protocol modified as in Govindarajulu et al. (2012) and PCR amplification was performed with 1X standard reaction buffer (New England Biolabs), 100uM of each dNTP, 0.5uM of each forward and reverse primer, 1.5 units of Standard Taq polymerase (New England Biolabs) and 1.5 ul of genomic DNA in a 20 ul reaction. The amplification conditions were: 2.5 minutes at 95°C, followed by 35 cycles of 95°C for 30 seconds, 55 and 56°C for 30 seconds, and 72°C for 60 seconds, and a final extension at 72°C for eight minutes (Table S2). The amplified products were Sanger sequenced and aligned using Sequencher ver 4.8 (Gene Codes Corp, Ann Arbor, Michigan) and SNPs were scored. For all crosses, we analyzed all segregating SNPs observed within each sequenced PCR product, whether or not they were the same segregating SNPs observed in the linkage maps. Association between segregating markers and sexual phenotypes was assessed in the progeny sets using Fisher’s exact tests.

We identified potential candidate genes in the region matching male sterility using the Hybrid GeneMark Predictions (Shulaev et al. 2010) available from GDR: Genome Database for Rosaceae (www.rosaceae.org/species/fragaria/fragaria_vesca/genome_v1.1) supplemented with four previously unrecognized genes (Darwish et al. 2015). For functional annotation we relied on PLAZA 3.0 (Proost et al. 2015). We examined levels of gene expression in a floral development transcriptome (Hollender et al. 2014), especially at stages 8-10 which are important in pollen development (Hollender et al. 2012). To test for PPR enrichment in our genomic region of interest, we identified 653 genes annotated as “Pentatricopeptide repeat-containing protein” (PPR) and counted them in 1Mb nonoverlapping windows across the genome. We also examined the genomic location of the 98 PPR genes that have a putative ortholog (as determined by PLAZA 3.0) with the 26 fertility restorer-like (RFL) genes in *Arabidopsis thaliana* (Fujii et al. 2011).

## RESULTS

### Novel sex determining region discovered

Of the 53 total selfed progeny of NM-LNF23, 9 were female and 44 were hermaphrodite (1:3 ratio; χ^2^ = 2.7; *P* = 0.10; Table 1A). Of the 41 offspring that were part of the mapping population, 8 were female and 33 were hermaphrodite (1:3 ratio; χ^2^ = 0.07; *P* = 0.79). Using these data we unambiguously mapped male sterility in these plants to a region near the 3´ end of chromosome LG6 (Figure 1). Specifically, at 14 sites on 10 targeted sequence capture probes between Fvb6_34958975 and Fvb6_36048692 (Figure 2A; Table 2 I), we observed a perfect match to male function. These perfectly matching sites include the region Fvb6_35142k (also Sanger genotyped in other crosses; Table 2 II-A), at which two SNPs (Fvb6_35142280 = Sanger site 280; and Fvb6_ 35142453 = Sanger site 453) cosegregated perfectly with sex type (*P* = 0.0001). At these two sites, all 8 females are homozygous for one of the two parental haplotypes (“G_T” at the Fvb6_35142k sites; Table 2 I), while all 33 hermaphrodites were either heterozygous or homozygous for the other parental haplotype (“T_C or “G/T_C/T” at the Fvb6_35142k sites), consistent with recessive male sterility (LOD = 8.8; Figure 1). From this we infer the genotype of NM-LNF23 to be *Rr* at the male function locus and its female progeny to be *rr* and hermaphrodite progeny to be *RR* or *Rr* (Table 3). Adjacent to these markers, the nearest mismatching markers are upstream at Fvb6_34839229 and downstream at Fvb6_36607138, and thus the male sterility locus must occur in the 1.769Mb region between these two markers (Figure 2A). The SNPs just outside this region, including in regions Fvb6_34763k and Fvb6_36607k (Table 2 II-A) also segregated significantly with sex type (*P <* 0.001), although at Fvb6_34839229 and farther upstream, a single female mismatched, while at Fvb6_36607138 and father downstream, one females and two hermaphrodites mismatched.

**Figure 1.**
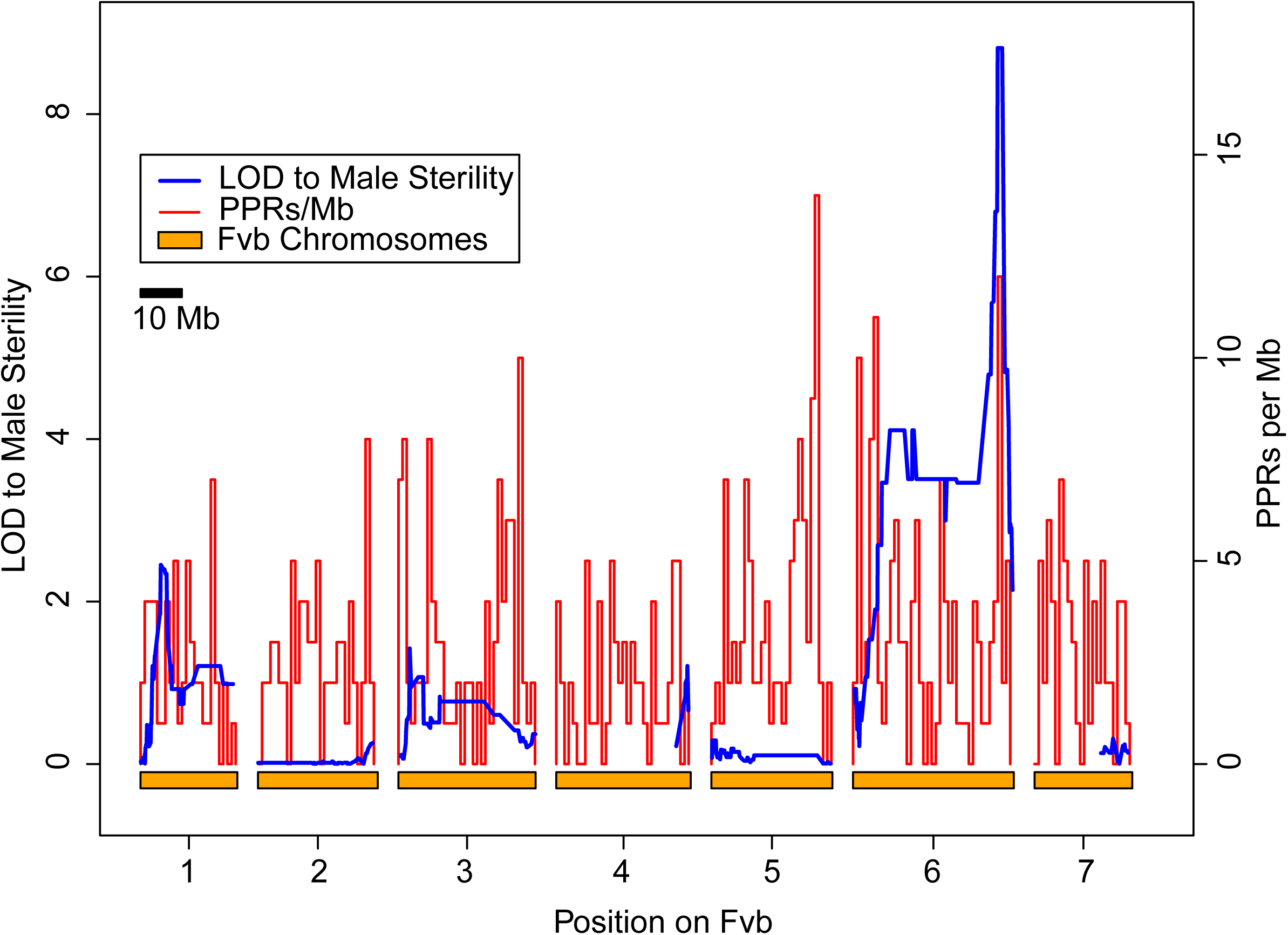
Map of male sterility in genome of hermaphrodite *F. vesca* subsp. *bracteata* from New Mexico (NM-LNF23).The seven chromosomes based on the Fvb reference genome (Tennessen et al. 2014) are denoted by orange bars along the *x*-axis and LOD scores associated with male function (blue line; left hand *y* – axis) and PPR gene density (PPR/Mb, red line; right hand *y* – axis) on the *y* – axes. A significant LOD score (>3) only occurs on LG 6, peaking at 8.8 for markers between 35.0-36.0 Mb. This region overlaps one of the two densest clusters of PPR genes in the genome, the other occurring on LG5.

**Figure 2.**
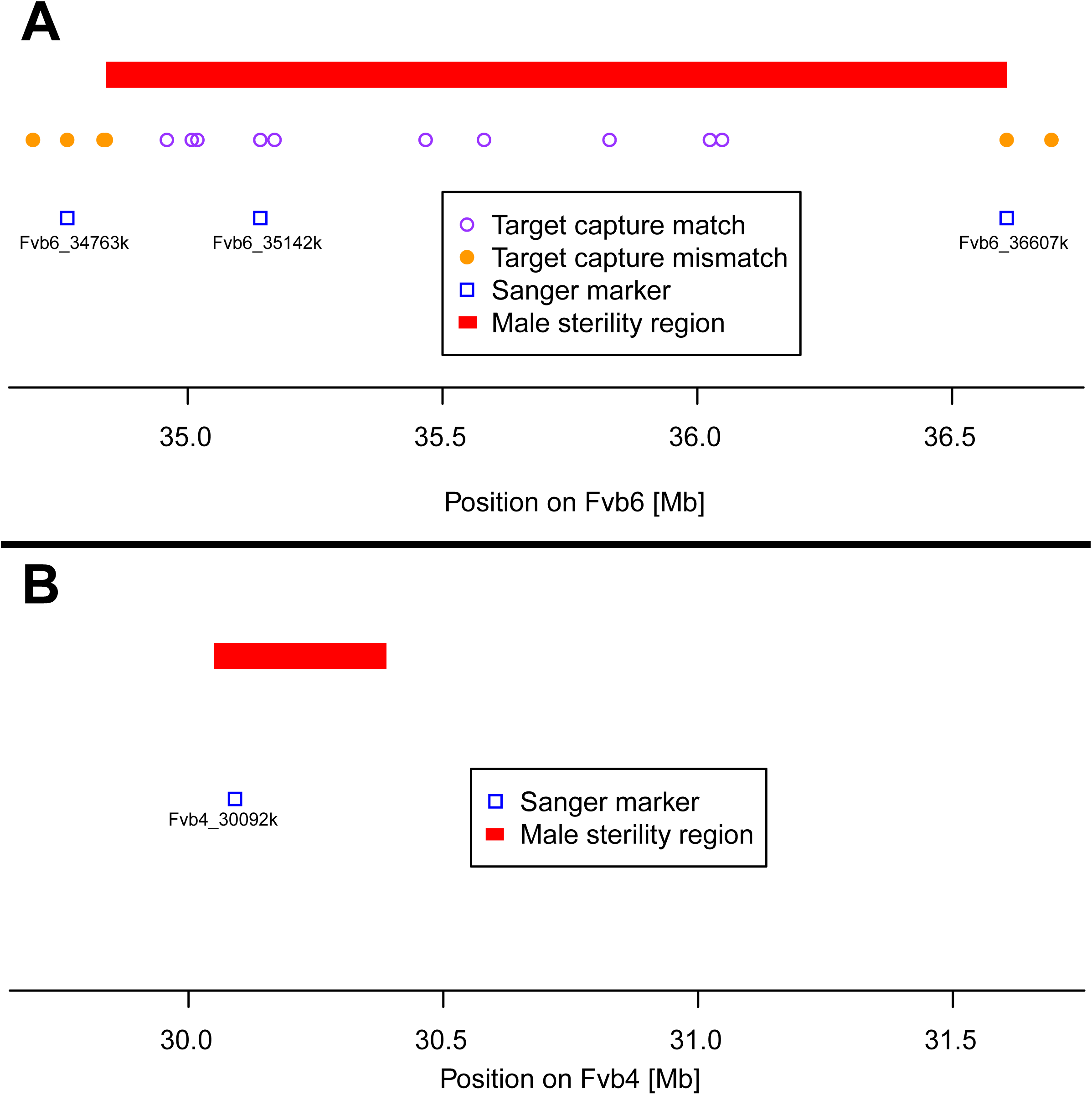
Male sterility genomic regions. (A). The LG6 locus (*R/r*). The male sterility region is defined as the span including 10 targeted sequence markers that perfectly match male sterility (see text). Three markers that were genotyped to confirm location and explore segregation with male sterility in other crosses with Sanger sequencing are noted by blue boxes (Fvb6_34763k, Fvb6_35142k and Fvb6_36607k). (B). The LG4 (*MS/mf*) locus. The male sterility region was mapped in Tennessen et al. (2013). Locations of Sanger markers used in this study are indicated by a blue box (Fvb4_30092k).

**Table 3.** Inferred genotypes of parents at sex determining loci in two populations of gynodioeicous *Fragaria vesca* subsp. *bracteata*. Plant identity, phenotypic sex (F = female; H = hermaphrodite) and putative genotype at loci mapped to linkage group 4 and 6 (LG4 717 and LG6), or unmapped but inferred from progeny segregation ratios (Locus *x*). At LG4 the *MS* allele codes for male sterility and is 718 dominant to mf which confers male fertility. At the LG6 locus, the *R* ‘restores’ male fertility and is dominant to *r* which does not (and 719 thus, codes for male sterility). At the unmapped locus *x*, the *T* codes male fertility and is dominant to *t* which codes for male sterility.

| Population | Plant ID | Sex | Inferred genotypes |  |  |
| --- | --- | --- | --- | --- | --- |
|  |  |  | Locus LG4 | Locus LG6 | Locus *x* |
| OR-MRD | OR-MRD93 | H | *mfmf* | *Rr* | *TT* |
|  | OR-MRD61 | H | *mfmf* | *RR* | *TT* |
|  | OR-MRD45 | H | *mfmf* | *RR* | *Tt* |
|  | OR-MRD30 | F | *MSmf* | *RR* | *TT* |
|  | OR-MRD27 | F | *MSmf* | *RR* | *TT* |
|  | OR-MRD90 | F | *MSmf* | *RR* | *TT* |
| NM-LNF | NM-LNF23 | H | *mfmf* | *Rr* | *TT* |
|  | NM-LNF25 | H | *mfmf* | *Rr* | *TT* |
|  | NM-LNF14 | H | *mfmf* | *Rr* | *Tt* |
|  | NM-LNF2 | F | *mfmf* | *rr* | *TT* |
|  | NM-LNF4 | F | *mfmf* | *rr* | *TT* |
|  | NM-LNF26 | F | *mfmf* | *rr* | *TT* |

The 1.768Mb region on Fvb6 that shows a perfect match to male-sterility contains 361 genes in the *Fragaria vesca* Hawaii 4 reference genome (Table S3). Including 182 that are upregulated in anthers, pollen or microspores, with one male gametophyte-specific gene, and several F-box proteins that have been seen to be upregulated in meiotic anthers at stage 9 (Hollender et al. 2014). Other gene classes showing expression changes in developing anthers included Kelch repeat and Leucine-Rich Repeat (LRR) proteins. This region also includes 15 PPR genes (Table S3), 12 of which occur in the 1Mb span between Fvb6_35Mb and Fvb6_36Mb, and 10 of which occur in the 0.5Mb span between Fvb6_35Mb and Fvb6_35.5Mb. This cluster of PPRs is unusually dense in this region relative to the rest of the genome (mean genomic PPR density = 2.9 per Mb). In fact, only one other genomic location, on Fvb5, contains a higher density cluster of PPRs (Figure 1). Although none of these PPRs are orthologs of the *Raphanus Rfo* fertility restorer and *Arabidopsis* fertility restorer-like (RFL) genes (Fujii et al. 2011), four are in the PLAZA gene family HOM03D000002, and one of these (gene04450) is predicted to be mitochondrial targeted (Table S4). This gene family contains the *Raphanus Rfo* gene and 25 of the 26 *Arabidopsis* RFL genes, as well as PPRs at two fertility-restorer loci recently identified in a hybrid cross almond×peach (Donoso et al. 2015). The RF1 locus on peach LG2 has 1 of 4 PPRs in this gene family, while the RF2 locus on peach LG6 has a dense cluster of 12 HOM03D000002 PPRs in 843.5 kbp. Five of these are considered orthologs of the *Arabidopsis* RFL genes (Table S4).

### Sex expression in additional NM-LNF crosses and confirmation of LG6 region of influence

The progeny of selfed hermaphrodite NM-LNF25 showed a 1:3 female to hermaphrodite sex ratio (χ^2^ =1.42; *P* =0.23) which is consistent with heterozygosity at a male function locus (*Rr*) (Table 1A; Table 3). The progeny of selfed hermaphrodite NM-LNF14, however, deviated significantly from a 1:3 female to hermaphrodite sex ratio (χ^2^ =10.00; *P* =0.002); leading us to propose a third, yet unmapped, locus affecting male function (locus LG*x*). The progeny sex ratio from selfed hermaphrodite NM-LNF14 is consistent with a 9:7 sex ratio (χ^2^ =0.47; *P* =0.49) that could result from selfing of a plant heterozygous at two male function loci (*Rr* and *Tt*) (Table 1A; Table 3). Sequencing the progeny from this cross for the Fvb6_35142k locus and evaluating the positions 35142280 and 35142453, we find that of the 15 hermaphrodite offspring, all either G, T or G/T, C/T at these two sites, but none of them are T_C. This makes sense because T (at 35142280) and C (at 35142453) are in coupling with *r* at the *R* locus, so all 7 T_C offspring are *rr* homozygotes and therefore female. The other female offspring are presumably *tt* at locus LG*x*. All other parents from NM-LNF are proposed to be *TT* at this locus (Table 3).

Crosses involving the NM-LNF hermaphrodites as sires each with three NM-LNF females produced progeny sex ratios (1:1) consistent with hypothesized genotypes of *Rr TT* for two of the hermaphrodites, and *Rr Tt* for one (NM-LNF14) and females all *rr TT* (all χ^2^ < 1.6, *P* > 0.19; Table 1A). The reciprocal crosses between NM-LNF25 and NM-LNF23 also produced progeny sex ratios (1:3) consistent with hypothesized genotypes of *Rr TT* for both of these hermaphrodites, though one of these crosses had very low seed set (*P* > 0.80; Table 1A; Table 3). The reciprocal crosses of NM-LNF14 with NM-LNF23 and NM-LNF25 also produced few seeds. The one cross that produced sufficient seeds NM-LNF14xLNF23 segregated in a manner consistent with the putative genotypes (1:3; *P* = 0.43).

We sequenced 20 progeny from female NM-LNF2 by hermaphrodite NM-LNF23 cross for the Fvb6_34763k and Fvb6_35142k markers and found three SNP that segregated with sexual phenotype (Table 2 II-A). At Fvb6_35142k positions 35142280 and 35142453 the *R* haplotype is T_C, and the *r* haplotype is G_T, all 9 hermaphrodites are T_C and 9 of the 11 females are G_T/T_C (*P* <0.0003). SNP position 34763440 of Fvb6_34763k also segregates with sex (*P*< 0.0001), and is consistent with only the same two female types mismatching, although the genotype is missing for one putatively mismatched female at Fvb6_34763k (Table 2 II-A). Although it would be desirable to genotype progeny in the Fvb6 34.8-36.6 Mbp region from the NM-LNF25 crosses, this individual was not heterozygous for any of our SNP markers in this region (see Table 2 I).

In sum, there is a clear indication from the combined phenotypic and genotypic data that the Fvb6 34.8-36.6 Mbp region of the genome influences sex expression when either a female or a hermaphrodite from NM-LNF population is the maternal parent. However, there apparently is a third unmapped locus (LG*x*) influencing sex phenotype.

### Sex expression in OR-MRD crosses

To evaluate whether hermaphrodites from OR-MRD also carried sex determining loci we evaluated progeny sex ratios from three self-pollinated hermaphrodites. One (OR-MRD93) showed a pattern consistent with a 1:3 sex ratio (χ^2^ <0.5; *P* >0.50), but not 0:1 (χ^2^ =50.6; *P* <0.001), one (OR-MRD61) deviated significantly from 1:3 (χ^2^ =14.3; *P* <0.001) but fit a 0:1 ratio (*P* >0.30), and one (OR-MRD45) fit both 1:3 and 0:1 equally well (both χ^2^ <0.5; *P* >0.30), (Table 1B). From this we inferred the genotype of OR-MRD93 as a *Rr TT* heterozygote and OR-MRD61 as a *RR TT* homozygote (Table 3). Crosses conducted between OR-MRD61 and the other two OR-MRD hermaphrodites produced nearly exclusively hermaphrodite progeny (there was one anomalous female; Table 1B), corroborating the inferred genotype of OR-MRD61 as *RR TT*. The other hermaphrodite by hermaphrodite crosses, however, did not produce any (OR-MRD93 × OR-MRD61) or many viable seeds (between OR-MRD93 and OR-MRD45) (Table 1C). And although too few seeds (4 and 7 hermaphrodite progeny) were produced from OR-MRD93 × OR-MRD45 reciprocal crosses to differentiate between 1:1 or 1:3 ratios, crosses between OR-MRD45 and NM-LNF plants (see discussion below; Table 1C,D) clarify the genotype of OR-MRD45 as *RR* at the LG6 locus. In addition, because the OR-MRD45 self-cross produced a few female progeny (Table 1B), we deduce the genotype of OR-MRD45 as *RR Tt* (Table 3).

To determine whether sex of progeny from these hermaphrodites is determined by the same region as in NM-LNF hermaphrodites we genotyped progeny from OR-MRD93self at the LG6 markers (Table 2 II-B). SNPs at position 34763379 on Fvb6_34763k and at position 35142492 on Fvb6_35142k both segregated perfectly with sexual phenotype (*P*<0.0001) with females being T_G and hermaphrodites being G/T-/A/G heterozygotes or G_A homozygotes. Thus, we have evidence here that the male sterility region on LG6 also influences sexual phenotype in the OR-MRD population when hermaphrodites are the dam. The fact that the same genomic region determines sex in both OR-MRD93 with A mitotype and NM-LNF with F mitotype (Table S1) suggests that the same *R* locus is responsible for sex phenotype in both cytoplasmic backgrounds. However, different PPR genes among the 15 at this locus could be the functional restorer in different cytoplasmic backgrounds.

When all three OR-MRD hermaphrodites were used as sires on three different females from OR-MRD, all produced 1:1 female to hermaphrodite ratios (all χ^2^ <2.63; *P* >0.11; Table 1B). Such patterns are consistent with the OR-MRD30×OR-MRD60 cross mapped in Tennessen et al. (2013) where a dominant allele in coupling with male sterility on the LG4 chromosome determines sex. Females were thus inferred to be *MSmf* whereas hermaphrodites to be *mfmf* at this locus (Table 3).

Finally, genotyping the hermaphrodite progeny from the OR-MRD61 × OR-MRD93 cross showed no SNP segregating with sexual phenotype as expected from a *Rr* × *RR* cross (Table 2 II-B; Table 3). It also suggests that these sex determiners are allelic and function both on the A mitotype and C mitotype cytoplasm (OR-MRD93 and OR-MRD61, respectively) in the absence of the influence of *MS* at the LG4 locus.

### Inter-population crosses to evaluate interactions between LG4 and LG6 sex determining regions

To evaluate whether maternal control of sex at LG4 in OR-MRD extended to NM-LNF sires we performed inter-population crosses on all three OR-MRD females. Eight out of the nine crosses produced 1:1 sex ratios (χ^2^ <2.57; *P* >0.22); the remaining cross (OR-MRD27*×*NM-LNF23) was skewed toward females (χ^2^ =4.77; *P=*0.03) (Table 1C).

For three of these crosses we genotyped progeny for LG6 and LG4 markers (Table 2 II-C). None of the segregating SNPs on LG6 segregated with sexual phenotype in any cross (*P* =0.67-1), but all three had SNPs at LG4 Fvb4_30092k positions 30092575 and 30092582 segregating with sex in the progeny (*P* <0.0001). All female progeny were heterozygotes G/C_A/C whereas hermaphrodites were G_G homozygotes in families of OR-MRD30*×*NM-LNF23 and OR-MRD90*×*NM-LNF23 (Table 2 II-C). In the OR-MRD30*×*NM-LNF14 family all female progeny were A homozygotes whereas hermaphrodites were A/C heterozygotes at the LG4 Fvb4_30092k position 30092575. Taken together these results are consistent with epistasic dominance of the *MS* allele at the LG4 locus over the *R* allele at the LG6 locus in determining sexual phenotype when the maternal parent is a female from OR-MRD.

To evaluate whether the sex-determiners at the LG6 locus of hermaphrodites from OR-MRD and NM-LNF are allelic (e.g., alleles at the same locus) and determine sex in the absence of the *MS* allele, we assessed progeny from OR-MRD hermaphrodite dams with NM-LNF hermaphrodite sires (Table 1C). No seed was produced when NM-LNF14 was a father, but crosses with either NM-LNF23 and NM-LNF25 as a sire produced progeny sex ratios that conformed to expectations (χ^2^ <2.6; *P* >0.22 Table 1C) based on inferred genotypes (Table 3). These also provide additional support for OR-MRD45 as *RR* (and not *Rr*, as all 3 crosses deviated significantly from 1:1 *P* <0.0001).

To follow up on these phenotypic findings, we genotyped the progeny from OR-MRD93 and OR-MRD61when pollinated by NM-LNF23. In the first cross, all four SNPs on Fvb_35142k segregated significantly with sex type (*P* =0.0002-0.05; Table 2 IIC), indicating that *R* alleles in both parents influence male fertility. In the second cross, as expected if OR-MRD61 is *RR*, none of the LG6 SNPs segregated with sex and all progeny are hermaphrodite.

To further explore the interaction between LG6 regions we assessed inter-population crosses between NM-LNF females and OR-MRD hermaphrodites (Table 1C). When crossed to OR-MRD45 and OR-MRD61, all three LNF females produced 0:1 sex ratio in the progeny as expected if the dam is *rr* and both the sires are *RR* at LG6 and *mfmf* at LG 4 (Table 3). The exception, however, was that when OR-MRD93 was the sire all three NM-LNF females produced sex ratios that deviated significantly from the 1:1 expectation (all *P* < 0.01; Table 1C). In fact, for all three crosses the female: hermaphrodite ratios fit a 2:1 (all *P* >0.80), potentially indicating a lethal genotype. To explore this hypothesis we genotyped the progeny from the NM-LNF2×OR-MRD93 cross and found the LG6 SNPs at position 34763379 on Fvb6_34763k and at position 35142492 on Fvb6_35142k both segregated perfectly with sexual phenotype (*P*<0.0001), females T_G and hermaphrodites G/T_A/G as predicted by the identified sex locus. Moreover, we recover all four expected genotypes in the offspring (consider both the maternally segregating genotypes (e.g. at Fvb6_34763308), and paternally segregating genotypes (e.g. at Fvb6_34763379): G/T_T, T_T, G/T_G/T, and T_G/T). Thus, the data are consistent with an unlinked locus causing 25% of offspring who are also *Rr* hermaphrodites to die. Inter-population crosses between NM-LNF hermaphrodites as dams and OR-MRD hermaphrodites as sires did not produce many seeds (Table 1C). However, the fact that the cross between NM-LNF14 and OR-MRD45 produced two females out of four total progeny is consistent their inferred heterozygous genotypes at the unmapped locus *x* (e.g. *mfmf Rr Tt* and *mfmf RR Tt*, respectively; Table 3).

## DISCUSSION

The work presented here combined with evidence from Tennessen et al. (2013), Stanley et al. (2015) and Govindarajulu et al. (2015), represents the first genomic evidence of both CMS and nuclear genes (fertility restorers) in a wild gynodioecious species. Moreover, demonstration of population variation in sex determiners provides insight into the potential complexity of cyto-nuclear gynodioecy in the wild and provides a road map for new hypotheses for the evolution of sex chromosomes in the related octoploid *Fragaria.*

### A multilocus model of sex expression in *F. vesca* subsp. *bracteata*

We have genetically mapped two unlinked epistatically interacting loci that determine sexual phenotype in wild populations of gynodioecious *F. vesca* subsp. *bracteata*. A single dominant *R* allele at the novel locus fine-mapped on LG6 restores fertility in the absence of *MS* allele at the LG4 locus previously identified by Tennessen et al. (2013). This newly identified LG6 34.8-36.6 Mbp region coincides with the second most dense cluster of PPR in the *F. vesca* reference genome and houses 361 genes. Four of the 15 PPRs (Table S3) can be considered top candidates for the fertility restorer, due to their presence in the HOM03D000002 gene family that contains the radish *Rfo* gene (the only confirmed CMS-interacting fertility-restorer in the rosid clade, Chen and Liu 2014), *Arabidopsis* RFL genes (Fujii et al. 2011) and putative RF loci in peach (Donoso et al. 2015). One of the four *F. vesca* genes is predicted to be mitochondrial targeted (Proost et al. 2015) and the other three do not have an organellar prediction. Other genes potentially involved in male function (upregulated in meiotic anthers; Hollender et al. 2014) also reside in this region, although not more so than the rest of the genome. Nonetheless it is intriguing that several important gene domains (F-box, Kelch repeat and LRR) occur here and in the LG4 male sterility region (*MSmf*) (Tennessen et al. 2013). Additionally, the action of miRNAs have been proposed as a route to sex determination (Akagi et al. 2014; Fagegaltier et al. 2014), and seven genes that are miRNA targets reside in the LG6 region (Table S3). Only one of these is a PPR, but it is cp-targeted and not in the HOM03D000002 gene family. Furthermore, none of these seven genes are targets for the novel F-box associated *Fragaria* miRNA family recently described in *F. vesca* (Xia et al. 2015).

A third locus (LG*x*) is also present and segregation analysis suggests it is unlinked to the LG6 locus (i.e., 9:7 ratio in the progeny of LNF14self, Table 1,2). Extensive mapping of genome-derived sequences around the sex-determining regions in several crosses revealed that the *MS* allele was epistatically dominant to *R* at LG6 and *T* at LG*x*. In fact, segregation results suggest that all loci interact, but functional studies will be needed to determine if and how they interact molecularly and biochemically.

Toward this end, we propose a conceptual model of these gene interactions (Figure 3) that draws on evidence from the current study and Tennessen et al. (2013) along with knowledge of the presence of a CMS-like *atp8-orf225* in the mitochondria (Table S1; Stanley et al. 2015; Govidarajulu et al. 2015). We hypothesize that a CMS locus that disrupts pollen development by production of a toxic protein or energy deficiencies (reviewed by Chen and Liu 2014) exists ancestrally. Male function is then restored by one or more copies of *R* at the LG6 34.8-36.6 Mbp locus. The presence of a cluster of PPRs at this locus, and the dominance of the restoring allele are consistent with data and theory of restorers of CMS (Delph et al. 2007; Chen and Liu 2014). The LG4 locus, which is not associated with annotated PPRs (Tennessen et al. 2013) is an inhibitor and a single dominant allele *MS* is sufficient to block the action of the LG6 restorer. Although of unknown function the LG4 locus may be a necessary cofactor required by the PPR or another mitochondrial sorting gene involved in nuclear-mitochondrial cross-talk (figure 3 in Chen and Liu 2014). The *MS* allele is both dominant to *mf* and epistatically dominant to *R.* Homozygosity for the *t* allele at the third unmapped locus on LG*x* also leads to disruption of pollen production, and this locus could represent an independently evolved restorer (*T* allele), as cyto-nuclear theory predicts successive ‘waves’ of restoration (reviewed in Deph et al. 2007), and multiple restorer systems are common, with some loci at or near fixation (e.g., Garraud et al. 2011; Caruso and Case 2013). This is especially true when there is polymorphism in CMS (e.g., Dudle et al. 2001; Garraud et al. 2011) as there appears to be in *F. vesca* subsp. *bracteata*. (Table S1; Stanley et al. 2015). If so, the products of the LG4 locus might also be able to block its action. Alternatively, the LG*x* locus could be a novel recessive male sterility locus that acts independently of the others (as seen in Irkaeva et al. 1993). By this same token, the LG6 locus may also be a recessive male sterility locus that interacts with products of the LG4 locus unrelated to CMS, but this possibly seems less likely given the general predominance of multiple nuclear-cytoplasmic interactions in gynodioecy and the specific evidence of both PPRs and a CMS-like *atp8-orf225* in gynodioecious *F. vesca* subsp. *bracteata*.

**Figure 3.**
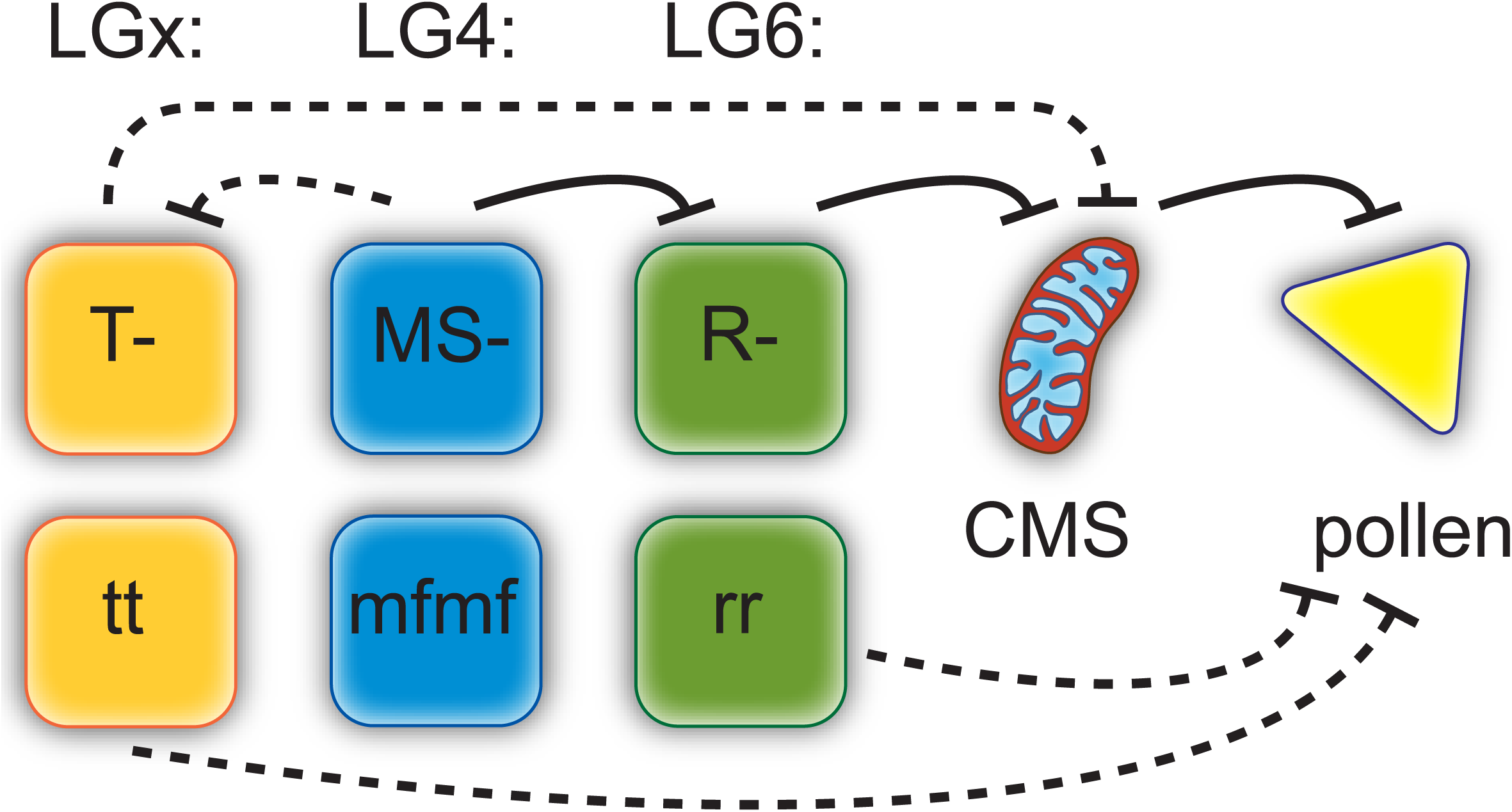
Conceptual model of male sterility loci in *F. vesca* subsp. *bracteata*. Black lines represent potential inhibitory interactions between genes. Dashed lines are uncharacterized genetically or posed as alternatives to the solid lines (see text). Three nuclear loci are represented by their known (LG4 and LG6) or unknown (LG*x*) location in the genome. The hypothesized involvement of these loci with a cytoplasmic male sterility (CMS) locus that blocks pollen development. The LG6 locus (*R/r*) is a restorer locus at which one copy of a dominant allele is sufficient to block CMS and restore fertility; thus male sterility is recessive at this locus. The LG4 locus (*MS/mf*) is an inhibitor at which a single dominant allele is sufficient to block the LG6 restorer. Thus the *MS* allele is both dominant to *mf* and epistatically dominant to *R.* Homozygosity for the *t* allele at the third unmapped locus on LG*x* also leads to disruption of pollen production, or could act as a restorer (*T* allele) similar to the LG6 locus.

The layers of interaction in this model may seem complex, but likely still reflect a simplification of the web of genetic mechanisms involved in cyto-nuclear gynodioecy in the wild. Much of what we know about these cyto-nuclear interactions comes from hybrids of hermaphroditic wild species (e.g., *Mimulus guttattus*, Barr and Fishman 2011; *Arabidopsis lyrata,* Aalto et al. 2013) or crop species where restorers are studied in fixed genetic backgrounds (reviewed in Chen and Liu 2014). In fact it has been proposed that one advantage of crop systems for understanding elements of cyto-nuclear gynodioecy is the reduced variation (Delph et al. 2007). However, these simplified systems may not represent the reality of sex determination in wild gynodioecious species which are subject to multiple and varied forces of drift, selection and gene flow. Our genomic evidence of both cytoplasmic and nuclear players join a small number of studies that have genetically mapped nuclear sex-determining loci in wild gynodioecious plants (Touzet et al. 2004), or have performed extensive crosses to demonstrate the existence of both nuclear and cytoplasmic contributors (e.g., Van Damme et al. 2004, Garraud et al. 2011). The next steps will involve functionally verifying the genetic interactions. Several types of approaches will be required for this and to fully test the working model we have proposed--including finer mapping of nuclear sex loci and transformation of both candidate nuclear restorer genes and the CMS-like *atp8-orf225* in the mitochondrial genome (see Chen and Liu 2014).

### Evolutionary inferences from population differentiation in sex determination

We observed population variation in the frequency of sex determiners, leading especially to differences in the genotypic composition of females. All examined females in OR-MRD have dominant male sterility (*MSmf R_ TT* genotypes), whereas all the females in NM-LNF have only recessive male sterility (*mfmf rr TT* genotypes). It should be noted that *mfmf rr TT* females are possible from selfed hermaphrodites in OR-MRD (i.e., OR-MRD93), but would be expected to be rare given the low frequency of *r* in this population. More generally, if our sampling is representative of the frequency of genotypes in the populations, then *r* is more frequent (1:11 vs. 9:3; *P* < 0.01) in NM-LNF than OR-MRD, while *MS* is more frequent in OR-MRD than NM-LNF, though the latter is not statistically significant (3:9 vs. 0:12; *P* > 0.05). This spatial genetic structure of sex determiners could provide insights into past evolutionary dynamics. Cyto-nuclear coevolution can lead to multiple restorer loci and polymorphism at both nuclear and mitochondrial genes (e.g., Garruad et al. 2011), and local processes are expected to result in spatial structure for restorer and CMS haplotypes (Bailey and McCauley 2005). So it is especially intriguing that the nuclear differences mirror the strong population differentiation in the mitotypes between the study populations. The F mitotype is only found in NM-LNF, whereas the B and C mitotypes found in OR-MRD are more widespread (Stanley et al. 2015). If, as our results suggest, LG6 and LG*x* are restorer loci then *R* and *T* can restore all three mitotypes, and this would imply a greater spatial range for restorers than mitotypes. Such a pattern has been inferred from crosses with differing cytotypes in *Plantago coronopus* (Van Damme et al. 2004) and *Silene nutans* (Garruad et al. 2011). In contrast, the dominant *MS* allele at LG4 appears restricted to the northwest (OR-MRD, in this study and Tennessen et al. 2014; and possibly ‘HP’ population in northern California, Ahmadi and Bringhurst 1991), perhaps indicating that it evolved after the LG6 and LG*x* loci, consistent with our multi-layered hypothesis for sex determination (Figure 3). The interactions between these genes likely contribute to high variation in sex ratios across space in *F. vesca* subsp. *bracteata* (0-46% females; Stanley et al. 2015). More crosses between populations of varying distances across the range (e.g., Bailey and McCauley 2005) and/or targeted resequencing of the sex determining regions would shed light on the wider geographic context for the players in the *F. vesca* subsp. *bracteata* sex determination system and our sex-determination hypothesis. This combined with fitness consequences of specific genotypes is needed to test evolutionary hypotheses for maintenance of multiple restorers and mitotypes in this species (e.g., Caruso et al. 2012).

The genetic differentiation of NM-LNF and OR-MRD is also evident in Stanley et al. (2015) where STRUCTRE analysis based on nuclear markers distinguished OR-MRD as belonging to cluster 1 while NM-LNF to cluster 2 (more closely allied with subspecies *F. vesca* subsp. *americana*). Similar distinction was seen in the chloroplast. Such differentiation could explain not only the differences in prevailing sex determiners but also the asymmetry in success of the interpopulation HxH crosses. Crosses with a OR-MRD hermaphrodite as a sire on a NM-LNF hermaphrodite as a dam produced fewer seeds than the reciprocal, potentially reflecting cyto-nuclear incompatibilities (Rieseberg and Blackman 2010) that interact with the sex determiners because similar deficits were not see when females were the dams. Alternatively these could reflect exposure of costs associated with restoration that may be complex (Caruso et al. 2012). It is notable in this regard that NM-LNF14 performed poorly as a sire and a dam, and also was the only plant inferred to be heterozygous at both the putative fertility-restorer loci (Table 3).

### Relationship to sex determination octoploids and other Fragaria

The present results can be considered in the context of the known sex determining regions in the two octoploid descendants of *F. vesca* subsp. *bracteata*. In both *F.viginiana* and *F. chiloensis,* linked male and female sterility loci reside on a chromosome in homeologous group VI (LG VI-Av in *F. chiloensis,* and VI-B2 *F.viginiana;* Goldberg et al. 2010; Spigler et al. 2011; Tennessen et al. 2014). Thus, it is intriguing to identify a locus on LG6 in *F. vesca* subsp. *bracteata* that also affects sexual expression. The fact that the sex determining region in *F. vesca* subsp. *bracteata* is near the 3′ end of LG6, as is the sex determining region of *F. chiloensis* (Goldberg et al. 2010), raises the possibility they are homologous. However, the *F. vesca* subsp. *bracteata* LG6 locus (*Rr*) codes for recessive male sterility, whereas the LGVI-Av locus in *F. chiloensis* has dominant male sterility. This difference alone might suggest they are not the same locus, but it could also reflect a turnover in the sex determining chromosome. Heterogametic transitions are theoretically possible and can transition from male heterogamety to female heterogamety when release from mutational load is followed by sexually antagonistic selection (Blaser et al. 2014).

If, on the other hand, the *F. vesca* subsp. *bracteata* LG6 locus and the LG VI-Av sex determining locus in *F. chiloensis* are entirely unrelated, the discovery of the *F. vesca* subsp. *bracteata* LG6 locus provides additional evidence that LG6 is predisposed to be a sex determining chromosome, as initially proposed by Spigler et al. (2011) with respect to *F. viginiana* and *F. chiloensis*. Some autosomes are thought to be prone to hosting sex determining regions because they house many genes that can affect male function, or because a chromosome that already has been involved in sex determination is more likely to seize back this role in the future (Graves and Peichel 2010, Blaser et al. 2014). This could be facilitated by transposition of genes already involved in sex expression (e.g., Hughes et al. 2015), making the LG4 locus a candidate for such transposition during the origin of the octoploids producing a gene complex in *F. chiloensis*. Finer mapping and deep resequencing of the sex determining region in *F. chiloensis* is underway to test these ideas. In either event, the present results suggest great potential for sexual lability in *Fragaria,* as male sterility evolves frequently, independently and via different genetic mechanisms. Indeed, recent phylogenetic character state transition analysis detects frequent transitions to and from separate sexes (dioecy) in the *Fragaria* (Goldberg et al. unpublished). The present study allows us to speculate that this lability is facilitated by ancient evolution of CMS and repeated evolution of restorers and/or other suppressors of male function, and turnovers in sex determining chromosomes.

## ACKNOWLEDGEMENTS

The authors thank C. Kustek, L. Stanley, K. Schuller. H. Wipf for greenhouse, field or laboratory assistance, the Ashman lab for discussion that improved the manuscript. This work was supported by the University of Pittsburgh, the National Science Foundation (DEB 1020523 and RET/REU supplements to TLA and DEB 1020271 to AL), UPitt Mellon and NSF Predoctoral Fellowships to MK, and Norman H. Horowitz Fellowship, Pennsylvania Space Grant Consortium Research Scholarship and Fellowships funded by Howard Hughes Medical Institute to RD.

## Supplementary Files

Table S1. Cytotypes of *F. vesca* subsp. *bracteata* parents used in the crossing study. Population and plant ids and sex types are given along with sequence for chloroplast and mitochondrial genes. Cytotype codes follow Stanley et al. (2015).

Table S2 Primer sequences (forward/reverse), annealing temperatures and reference genome coordinates for eight informative polymorphic sites segregating with male sterility identified OR-MRD30xOR-MRD60 and NM-LNF23 self map cross population. Coordinates in FvH4 are with respect to the *F. vesca* ssp. *vesca* reference genome version 1. 0 (Shulev et al. 2010), while those in bold are on the scaffold scf0513158b (Tennessen et al. 2013). Coordinates in Fvb are with respect to *F. vesca* subsp. *bracteata* assembly of the reference genome version 1.0 (‘Fvb’ described in Tennessen et al. 2014).

Table S3. Genomic locations, functional annotations, and PLAZA 3.0 gene families of genes at the Fvb6 male sterility locus in *Fragaria vesca* subsp. *bracteata*. Notes: 1: Pentatricopeptide repeats (PPR), 2: the PPR gene family that contains known and suspected fertility restorers, 3: F-box proteins observed to be upregulated in meiotic anthers at stage 9, 4: miRNA target (Xia et al 2015), 5: unique to Darwish et al. (2015) annotation.

Table S4. Genomic locations and PLAZA 3.0 gene families of PPR genes at the peach RF1 and RF2 loci (Donoso et al. 2015). The PPR gene family that contains known and suspected fertility restorers denoted with * in the notes column.

